# Targeted capture of complete coding regions across divergent species

**DOI:** 10.1101/099325

**Authors:** Ryan K Schott, Bhawandeep Panesar, Daren C Card, Matthew Preston, Todd A Castoe, Belinda SW Chang

## Abstract

Despite continued advances in sequencing technologies, there is a need for methods that can efficiently sequence large numbers of genes from diverse species. One approach to accomplish this is targeted capture (hybrid enrichment). While these methods are well established for genome resequencing projects, cross-species capture strategies are still being developed and generally focus on the capture of conserved regions, rather than complete coding regions from specific genes of interest. The resulting data is thus useful for phylogenetic studies, but the wealth of comparative data that could be used for evolutionary and functional studies is lost. Here we design and implement a targeted capture method that enables recovery of complete coding regions across broad taxonomic scales. Capture probes were designed from multiple reference species and extensively tiled in order to facilitate cross-species capture. Using novel bioinformatics pipelines we were able to recover nearly all of the targeted genes with high completeness from species that were up to 200 myr divergent. Increased probe diversity and tiling for a subset of genes had a large positive effect on both recovery and completeness. The resulting data produced an accurate species tree, but importantly this same data can also be applied to studies of molecular evolution and function that will allow researchers to ask larger questions in broader phylogenetic contexts. Our method demonstrates the utility of cross-species approaches for the capture of full length coding sequences, and will substantially improve the ability for researchers to conduct large-scale comparative studies of molecular evolution and function.

## Introduction

It is difficult, in terms of the amount of resources needed, to study the evolution of a large number of complete genes from a large number of taxa, but continued advances in next-generation sequencing (NGS) technology have made this approach more feasible within reasonable time-frames and budgets. Despite these advances, sequencing entire genomes is generally too time-consuming, and too costly, on comparative taxonomic scales, and produces much more data than necessary for most evolutionary questions. PCR, on the other hand, still excels at sequencing small numbers of genes, but quickly becomes cost ineffective when large numbers of genes are required, while primer design and optimization becomes time inefficient across divergent species (Mamanova et al. 2010; Shen et al. 2013). As a result, there is a need for methods that can efficiently sequence a large set of genes of interest from a large number of species. Currently, such data are limited to the relatively small number of sequenced genomes and a growing number of transcriptomes. RNA-Seq (Wang et al. 2009) is becoming increasingly popular for comparative studies (e.g., Kunstner et al. 2010; Brousseau et al. 2014; Gallant et al. 2014; Gerstein et al. 2014; LoVerso and Cui 2015; Yang et al. 2015; Havird and Sloan 2016; Phillips et al. 2016; Wu et al. 2016), but has several downsides including a reliance on fresh tissue samples and variation in transcript expression levels. However, there are many methods that can be utilized to target, enrich, and capture specific sections of the genome (for reviews see Mamanova et al. 2010; Teer et al. 2010; Mertes et al. 2011). Targeted capture (also called targeted enrichment, hybrid enrichment, or sequence capture) is one of these methods that has been shown to perform well and is gaining popularity (Albert et al. 2007; Hodges et al. 2007; Okou et al. 2007; Porreca et al. 2007; Gnirke et al. 2009; Summerer et al. 2009; Mamanova et al. 2010; Nijman et al. 2010; Teer et al. 2010; Teer and Mullikin 2010; Kenny et al. 2011; Mason et al. 2011; Mertes et al. 2011; Bi et al. 2012; Bundock et al. 2012; Crawford et al. 2012; Cronn et al. 2012; Faircloth et al. 2012; Grover et al. 2012; Lemmon et al. 2012; McCormack et al. 2012; Rohland and Reich 2012; Li et al. 2013; Ilves and Lopez-Fernandez 2014; Penalba et al. 2014; Bragg et al. 2016; Portik et al. 2016).

Targeted capture is a method to selectively enrich the genome for particular regions of interest by using a set of DNA or RNA probes as bait (Gnirke et al. 2009). This can either be done on a microarray (Albert et al. 2007; Hodges et al. 2007; Okou et al. 2007) or in solution (Gnirke et al. 2009), but the principle is the same. The probes are designed to complement the region(s) of interest, whether a small section of the genome or the entire set of protein coding genes (the exome). The probes are then allowed to hybridize with a gDNA library that has been fragmented to produce inserts in the range of 200–700 bp. Inserts that fail to hybridize are washed away thus selectively enriching the genome for the regions of interest. Sequencing can than proceed normally, including multiplexing many samples to increase efficiency.

Hybrid enrichment was originally proposed, and has been most widely used, to capture and resequence the human exome (e.g., Albert et al. 2007; Hodges et al. 2007; Okou et al. 2007; Porreca et al. 2007; Gnirke et al. 2009), and has since been applied to whole exome sequencing in other model species for applications such as variant discovery and population genetics (for reviews see Warr et al. 2015; Jones and Good 2016). Applications of whole exome sequencing to related species show a decline in performance with even small amounts of divergence (Vallender 2011; Jin et al. 2012; Jones and Good 2016). Consequently, capture across divergent species requires modifications and tends to focus on more conserved sequences or a smaller targets. Several cross-species approaches have been developed and used both at broad taxonomic scales to capture highly conserved (e.g., Lemmon et al. 2012) and ultraconserved (e.g., Crawford et al. 2012; Faircloth et al. 2012; McCormack et al. 2012) regions, at narrow taxonomic scales to capture the mitochondrial genome (Mason et al. 2011), and at varying scales to capture individual exons and partial coding sequences (e.g., Bi et al. 2012; Li et al. 2013; Ilves and Lopez-Fernandez 2014; Penalba et al. 2014; Bragg et al. 2016; Hugall et al. 2016; Portik et al. 2016). The focus of each of these methods, however, is solely on producing data for phylogenetic studies. As a result, they use automated methods to select targets favorable for capture, rather than using complete coding regions from specific genes of interest. The resulting data is thus useful for phylogenetic studies, but the wealth of comparative data that could be used for molecular evolutionary and functional studies is lost.

Here we adapt and expand this method to selectively enrich complete coding regions from specific genes of interest across broad taxonomic scales. The focus on specific genes allows selection of sequences associated with aspects of organismal physiology that can be used to address various research questions. The capture of specific complete coding regions, however, presents unique experimental and computational challenges compared to capturing computationally selected conserved regions or exons. Regions that have high divergence, are predicted to have poor hybridization, or that are too short cannot simply be excluded as is typically done (e.g., Lemmon et al. 2012; Ilves and Lopez-Fernandez 2014; Hugall et al. 2016). The computational assembly of the data is also more complex as the individual exons need to be assembled into a continuous sequence while removing the intronic sequence that will be enriched alongside the targeted exons. To address these issues, we use a unique probe design strategy and develop novel bioinformatics pipelines for the assembly and analysis of complete coding regions. We evaluate the effects of different assembly algorithms and references, and compare our method to alternative approaches, namely whole genome sequencing (WGS) and RNA-Seq. The method we develop allows complete coding regions to be captured from a set of genes of interest, while maintaining the ability to capture across divergent species. The resulting data is still useful for phylogenetic analyses, but importantly can also be applied to studies of molecular evolution and function that will allow researchers to ask larger questions in a broader phylogenetic context.

## Results and Discussion

Here we adapt solution-based targeted capture following Gnirke et al. (2009), Lemmon et al. (2012), and Faircloth et al. (2012), to selectively sequence complete coding regions from specific genes of interest across divergent species. We took the unique approach of manually curating a set of genes of interest. This differs from the approach employed by other cross-species targeted capture methods, which target ultra (e.g., Faircloth et al. 2012) or highly (e.g., Lemmon et al. 2012) conserved regions, or individual exons with specific properties (e.g., Li et al. 2013; Ilves and Lopez-Fernandez 2014; Bragg et al. 2016; Hugall et al. 2016; Portik et al. 2016) compiled using purely computational means, and also provides benefits over alternative NGS strategies (Table 1). The focus of the method presented here is on sequencing the complete coding regions from genes of specific interest for more broad evolutionary applications, in addition to phylogenetic reconstruction. A total of 166 genes of interest were targeted composed of 1435 individual exons. These included visual, housekeeping, and phylogenetic marker genes (Table S1).

**Table 1.** Comparison of pros and cons of different high-throughput sequencing strategies.

| Method | Pros | Cons | References |
| --- | --- | --- | --- |
| WGS | Produces data that can be used for many different applications | Expensive. Assembly can be difficult and time consuming | Koepfli et al. (2015) |
| RNA-Seq | Produces data that can be used for many different applications. Also provides information on expression levels. | Need fresh tissue for RNA. Obtaining sequences depends on expression levels and tissue/temporal-specific expression | Wang et al. (2009) |
| PCR | Can sequence small numbers of loci (up to ~100) efficiently across divergent species. | Primer design can be difficult, time consuming, and needs conserved regions. Can give biased results. | Meyer et al. (2008); Bybee et al. (2011); Shen et al. (2013) |
| RRL/RAD-Seq | Can produce 1000s of loci for use in intraspecific and shallow phylogenetic studies. | Only useful at very shallow time scales. | Altshuler et al. (2000); Miller et al. (2007); Baird et al. (2008) |
| Targeted Capture (Hybrid Enrichment) | Can sequence 100s to 1000s loci from species at varying levels of divergence. | Initial probe design can be expensive/time consuming and may require genomic resources. | See below |
| Whole Exome Capture | Produces data that can be used for many different applications. | Only applicable with model organisms and their very close relatives. | Albert et al. (2007); Porreca et al. (2007) |
| Conserved Region Capture | Sequence 1000s of loci that can be used for phylogenetic studies at shallow to deep timescales. | Data only useful for phylogenetic studies. | Faircloth et al. (2012); Lemmon et al. (2012) |
| Exon Capture | Sequence 100s to 1000s of individual exons across low to moderate levels of divergence. | Data only useful for phylogenetic studies. Performance decreases sharply with divergence. | Li et al. (2013); Bragg et al. (2016); Portik et al. (2016) |
| Complete CDS Capture | Produces data that can be used for many different applications. Applicable across divergent species. Can target genes with physiological relevance. | Manual curation can be time consuming. Guided assembly can require additional references sequences for divergent species. | Current study |

Abbreviations—**WGS**, whole genomes sequencing; **RRL**, reduced representations library; **RAD**, restriction site associated DNA; **CDS**, coding sequence.

Probes were designed from the individual exons comprising the coding regions of each of these genes as shown in Figure 1. To facilitate capture of complete coding regions across divergent species probes were designed from multiple reference species following Lemmon et al. (2012). This resulted in an increased diversity of sequences comprising the probes targeting each sequence and may have allowed hybridization to occur in regions that were otherwise too divergent, or missing, in a specific reference (Fig. 1C). Probes were extensively tiled (10x) across the reference sequences, which similarly increased the diversity of probes sequences available for hybridization, and may have both compensated for hybridization issues with individual probes (e.g., secondary structure, GC content) and allowed capture across divergent regions (Fig. 1C). Exons that were shorter than the probe length of 120 bp could not be tiled and instead had to be padded with non-homologous sequence to increase the length to 120 bp (Fig. 1B). In total we targeted 3888 exons from the reference species, which resulted in 45,895 probes after tiling and boosting to normalize coverage (*DRYAD link tbd*).

**Figure 1.**
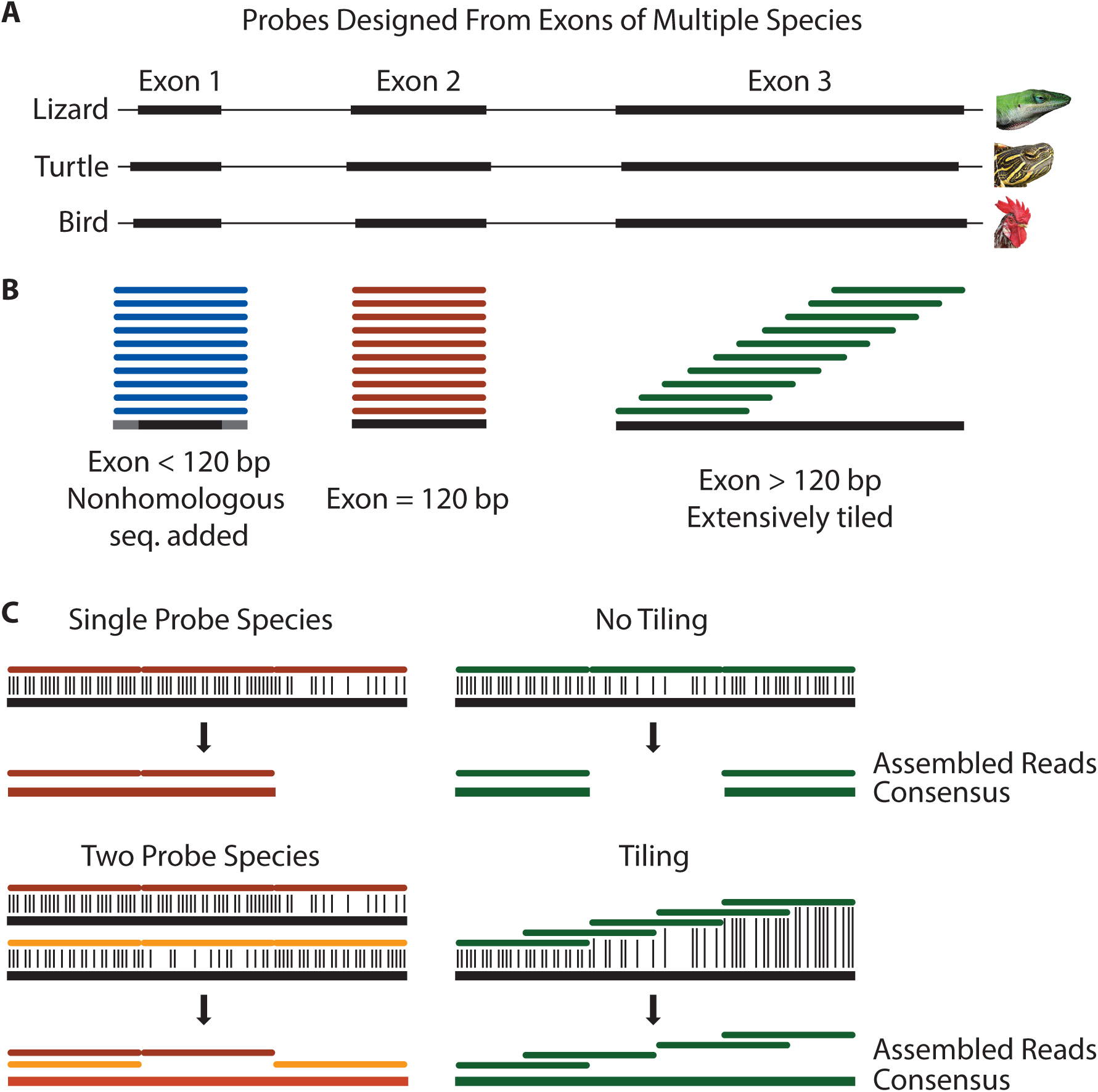
Cross-species hybrid capture methods. **A**, exons were extracted from the genomes of an average of three reference speces (anole, turtle, and chicken). **B**, probes were designed against each exon. Since probe length was constant at 120 bp, exons shorter than the probe length were padded with non-homologous sequence. Exons the same length as the probe matched exactly, while those longer were extensively tiled across the exon (10X coverage). The overall number of probes covering each base was normalized to ensure even coverage. **C**, multiple reference species and tiling were designed to help facilitate cross-species capture. For example, a region of high divergence may occur in one species and not another, or could still be captured by tiling across it.

As a proof-of-concept, we selected 16 squamate reptile (lizard and snake) species for hybrid enrichment and sequencing (Table S2). The species sampled here span a broad set of the major lineages of squamates, encompassing approximately 200 myr of divergence (Hedges et al. 2015). To recover the coding sequences, after hybrid enrichment and sequencing, we employed a guided assembly strategy utilizing custom assembly and analysis pipelines (*DRYAD link tbd*). Reads were assembled against a reference composed of the coding sequences of the targeted genes. The primary set of reference sequences were from *Anolis*, the probe species most closely related to the species we sequenced. Additional sequences were included from the other probe species for genes absent from the *Anolis* genome. Because assemblies were performed across species, we also used additional references compiled from snake genomic and transcriptomic data and from the *Gekko japonicus* genome, which were produced or became available after the design and synthesis of the probes. Several different assemblers were also used with different tolerances for mismatches. After assembly, consensus sequences were called and their identities confirmed by BLAST and, when necessary, phylogenetic analysis. We also calculated the completeness of each recovered sequence relative to the reference sequence (capture sensitivity).

Overall, the method was highly successful and nearly all genes were recovered (92% on average, Fig. 2, Tables S3-6). The level of recovery we achieved is higher than that of a previous cross-species capture study, which recovered from 16%–80% of coding sequence targets in comparisons that varied from having 106–299 myr of divergence (Li et al. 2013). Of the original 166 genes that we targeted, two genes (*ALB, SLC24A1*) were not recovered from any of the 16 squamate sequences (Table S4) or from any of the available squamate genomes *(Anolis carolinensis* (Alfoldi et al. 2011), *Python molurus bivittatus* (Castoe et al. 2013), *Ophiophagus hannah* (Vonk et al. 2013), *Thamnophis sirtalis* (Castoe et al. 2011), *Gekko japonicas* (Liu et al. 2015)) suggesting they were likely lost ancestrally in squamates. One gene *(CRYD2)* appears to be a lineage-specific duplication in some birds (e.g., chicken), based on its absence in the squamate (and other reptilian) genomes, and was also not recovered for any of the 16 species. The probe for one gene, *STRA6*. was later found to lack any homology with other *STRA6* sequences, which explained its lack of capture success. These four genes were thus excluded from further analysis. Three genes, *UBC, UBB*. and *UBI*, were found to all represent (at least part of) the same gene (which we term *UBC*) and thus were combined. This left a total of 160 for further analysis.

**Figure 2.**
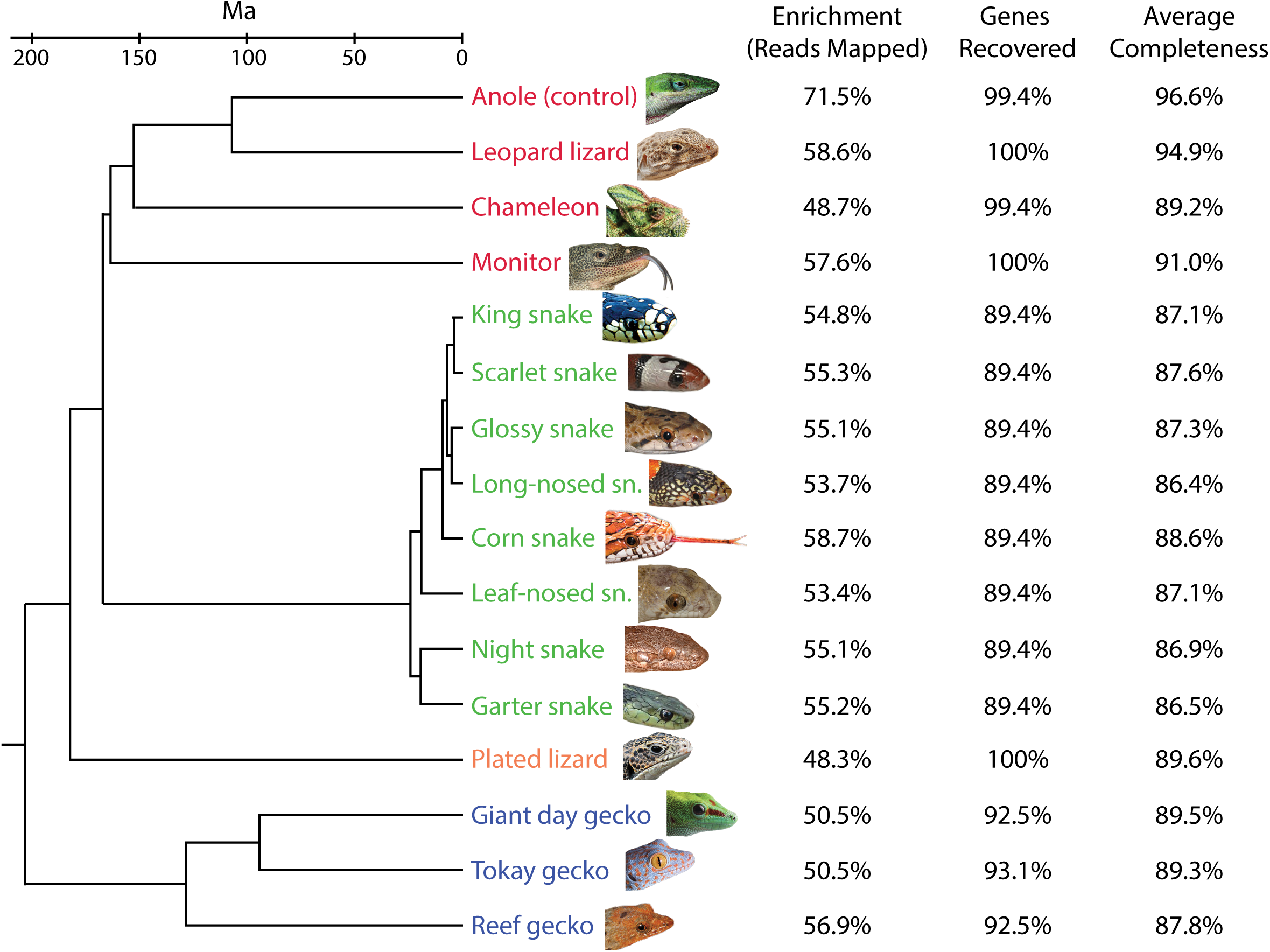
Species relationships of the 16 species sequenced and the enrichment, percent of genes recovered and the average completeness of those genes that were recovered. These results represent the combined best for the different assembly methods and references used. Genes were considered recovered if they were at least 5% complete and could be properly identified based on BLAST similarity and/or phylogenetic position. Species most closely related to the reference are shown in red, snakes in green, the plated lizard in orange, and geckos in blue. Tree topology based on Pyron et al. (2013). Divergence times from Hedges et al. (2015).

While the vast majority of genes were recovered in all species, a number of genes were not recovered in particular groups or individual species. In most cases, the lack of recovery appears to be due to gene loss rather than a failure of the method. For example, snakes and geckos are both known to have lost several visual genes (Zhang et al. 2006; Castoe et al. 2013). In colubrid snakes 17 genes were not recovered including the 10 opsin genes reported previously to have been lost in snakes *(NEUR2, NEUR3, OPN4m, parapinopsin, parietopsin, pinopsin, RH2, SWS2, TMT2, TMTa;* (Castoe et al. 2013)) as well as five lens crystallins (*CRYBA1, CRYBA4, CRYBB1, CRYBB3, CRYGN*), one phototransduction gene (*GRK1*), and one *HOX* gene *(HOXD12).* None of these genes were found in the *Python, Ophiophagus*, or *Thamnophis* genomes. Eleven genes were not recovered in the any of the three gecko species, including seven opsins (*NEUR2, NEUR3, OPN4m, parapinopsin, parietopsin, RH1, SWS2*), a lens crystallin (*CRYGN*), and three phototransduction genes (*CNGA1, PDE6B, PDE6G*). These genes were also absent from the *Gekko japonicus* genome. In *Phelsuma*, TMTa was also not recovered, and in *Sphaerodactylus CRYBA4* was not recovered. Both of these genes are present in the *Gekko* genome, so it is unclear whether they represent lineage-specific losses or a failure of the hybrid capture. All genes, other than those absent in all squamate taxa, were recovered for the other species, except for *CRYD* in *Anolis* and *pinopsin* in *Chamaeleo. CRYD* was also absent from the *Anolis* genome, although other genes absent from the *Anolis* genome were at least partially recovered.

Enrichment (or capture specificity) of the targeted genes was high, with an average (mean, throughout) of 55% of the reads mapping to the reference (Fig. 2, Table S3). This level of enrichment is much higher than that reported for other cross-species targeted capture methods (Bi et al. 2012; Lemmon et al. 2012; Ilves and Lopez-Fernandez 2014), which have reported mapping rates ranging from 5–33%. Our positive control, *Anolis*, had 71.5% of the reads mapping to the reference, which is close to the level seen in human resequencing studies, which achieve up to 80% reads on target (Mamanova et al. 2010). Coverage was also high with an average depth of coverage across all species and genes of 2159X that ranged from 2903X in *Anolis* to 1884X in *Phyllorhynchus* (Table S3). The high level of coverage we obtained suggests that we sequenced at much higher depth than will be necessary for future experiments. In subsequent experiments it would likely be possible to multiplex and sequence substantially more species, or many more genes, with the same amount of sequencing and still obtain high coverage.

Among the most important factors for the utility of the approach for wide evolutionary application is the completeness of the recovered coding regions (capture sensitivity). Completeness was generally high with an overall average of 89.1% using the best method and reference for each gene (Table S6). However, some genes were recovered with low completeness, all the way down to our cut-off of 5% (Table S4). Completeness varied considerably with the different assemblers and references used and varied among the genes and the species and will be discussed further below. While many cross-species capture studies do not report the completeness of recovered targets, Portik et al. (2016), who used transcriptome-based exon capture, reported an average completeness of 80% for their ingroup sample (up to 56 myr divergence) and 34% for their outgroup sample (up to 103 mry divergence), demonstrating a marked decrease in completeness with divergence not seen with our method.

### Reference sequences have a large effect on cross-species guided assembly

Three main sets of reference sequences were used in the assembly of the reads (*DRYAD link tbd*). The primary set of sequences was based on the coding sequences from *Anolis*, the same sequences used to develop the probes (*Anolis* reference). This set necessarily also included sequences from other taxa when a gene was missing or lost in *Anolis* (see Material and Methods for more details). When reads were assembled with BWA against this reference recovery and completeness were high for species more closely related to *Anolis*, but suffered for the more divergent species, especially the snakes and geckos (Table 2). To address this, we produced two more sets of reference sequences: one utilizing a *de novo* eye transcriptome from *Thamnophis sirtalis* and the *Python molurus bivittatus* (Castoe et al. 2013) and *Thamnophis sirtalis* genome assemblies (snake reference), and one with sequences from the *Gekko japonicus* genome (Liu et al. 2015) (*Gekko* reference). These references contained 139 and 110 sequences each as they only included snake or gecko sequences, respectively. When the colubrid snakes were assembled to the snake reference using BWA we obtained a small increase in the enrichment (percent of reads mapped to reference) and recovery of genes, but a ~20% increase in the completeness of the recovered genes present in both references (Table 2). A very similar increase in completeness (~19%) was found for the geckos when assembled to the *Gekko* reference (Table 3).

**Table 2.** Comparison of BWA assembly with the *Anolis* and Snake references.

| Species | Reference Sequences |  |  |  |  |  |  |  |  |
| --- | --- | --- | --- | --- | --- | --- | --- | --- | --- |
|  | *Anolis* (160 seqs.) |  |  | *Anolis* trim Snake (139 seqs.) |  |  | Snake (139 seqs.) |  |  |
|  | Enrich. | Rec. | Comp. | Enrich. | Rec. | Comp. | Enrich. | Rec. | Comp. |
| Anole (control) | 71.5% | 99.4% | 94.8% | 56.5% | 99.3% | 95.9% | 30.1% | 98.6% | 66.7% |
| Leopard lizard | 58.6% | 100% | 88.2% | 48.7% | 100% | 89.7% | 32.4% | 98.6% | 70.3% |
| Chameleon | 48.5% | 98.1% | 72.8% | 42.4% | 98.6% | 74.3% | 31.9% | 95.0% | 63.9% |
| Monitor | 56.7% | 100% | 77.7% | 48.6% | 100% | 79.8% | 39.6% | 97.8% | 71.5% |
| Glossy snake | 48.7% | 88.1% | 64.4% | 44.5% | 98.6% | 64.9% | 50.8% | 100% | 85.6% |
| Scarlet snake | 49.6% | 88.1% | 67.8% | 45.9% | 98.6% | 68.3% | 52.1% | 100% | 86.1% |
| King snake | 48.7% | 87.5% | 64.8% | 43.9% | 97.8% | 65.3% | 50.5% | 100% | 85.6% |
| Corn snake | 48.5% | 88.8% | 62.6% | 44.8% | 99.3% | 63.0% | 50.9% | 100% | 85.0% |
| Leaf-nosed sn. | 52.6% | 89.4% | 67.6% | 48.9% | 100% | 67.8% | 56.5% | 100% | 86.2% |
| Long-nosed sn. | 47.3% | 86.9% | 61.5% | 43.6% | 97.1% | 61.9% | 50.7% | 100% | 85.6% |
| Night snake | 49.1% | 87.5% | 64.2% | 45.8% | 97.8% | 64.7% | 52.4% | 100% | 85.0% |
| Garter snake | 49.1% | 86.3% | 63.8% | 45.8% | 96.4% | 64.2% | 51.6% | 100% | 84.7% |
| Plated lizard | 48.3% | 98.8% | 75.2% | 42.5% | 99.3% | 75.9% | 35.8% | 97.8% | 67.4% |
| Tokay gecko | 47.5% | 90.0% | 69.4% | 41.5% | 97.1% | 69.5% | 36.9% | 92.8% | 61.7% |
| Giant day gecko | 47.8% | 90.6% | 69.1% | 39.9% | 96.4% | 69.7% | 33.6% | 94.2% | 61.4% |
| Reef gecko | 53.4% | 90.6% | 65.6% | 46.7% | 97.1% | 66.1% | 38.7% | 93.5% | 58.7% |
| **AVERAGE** | **51.6%** | **91.9%** | **70.6%** | **45.6%** | **98.3%** | **71.3%** | **43.4%** | **98.0%** | **75.3%** |

To make the comparison between the *Anolis* and Snake reference even, the *Anolis* reference was trimmed to contain only the sequences present in the Snake reference (*Anolis* trim Snake). Enrichment was measured as the percent of reads mapping to the reference (uncorrected for genome size). Recovery was calculated as the percent of the 165 targeted genes that had at least 5% completeness and that could be identified based on BLAST similarity and/or phylogenetic analysis. Completeness was calculated relative to the reference sequence for those genes identified as recovered using the BWA assembly method and the best reference for each gene. Abbreviations—**Enrich.**, Enrichment; **Rec.**, Recovery; **Comp.**, Completeness; **sn.**, snake.

**Table 3.**
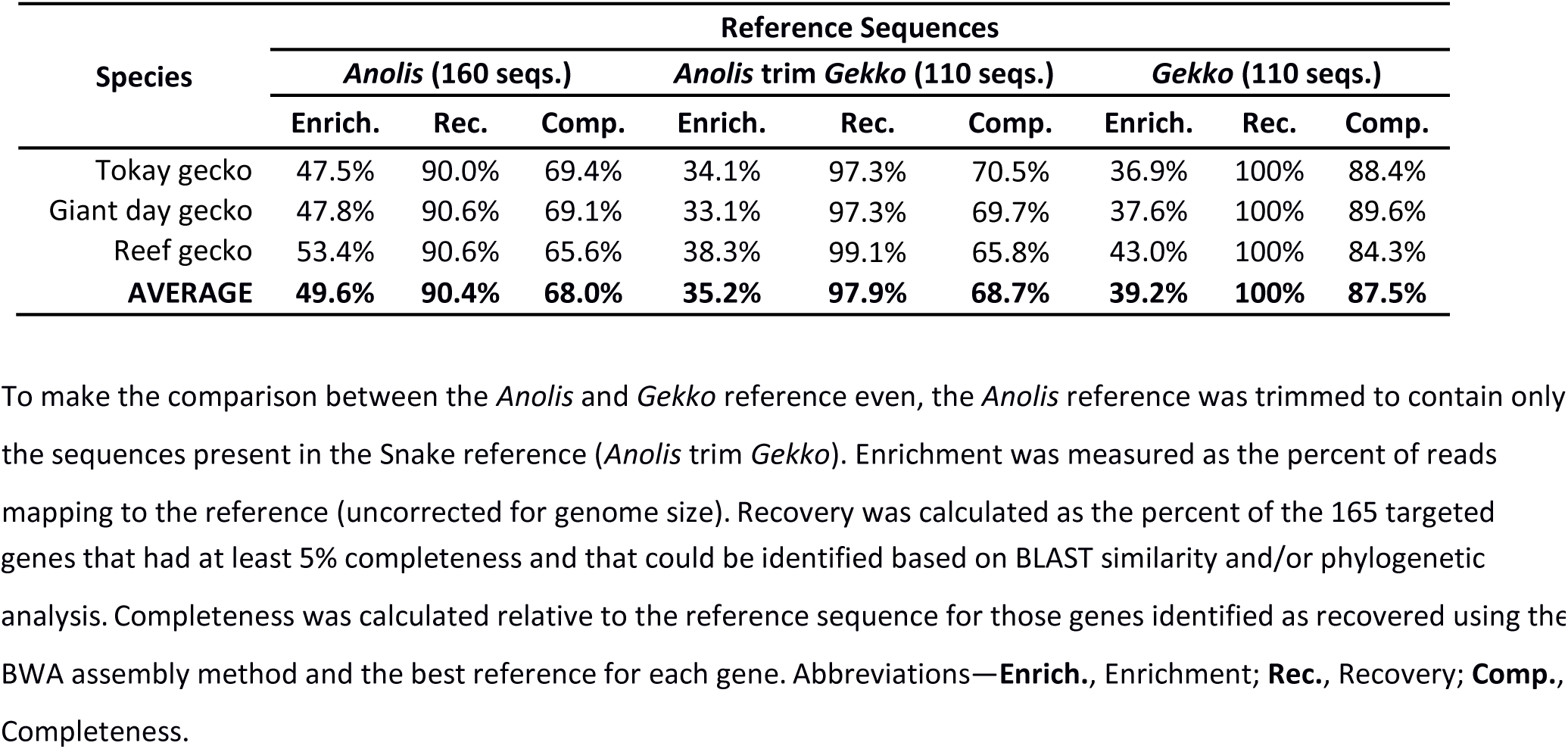
Comparison of BWA assembly with the *Anolis* and *Gekko* references.

The effect of using different reference sequences was also be demonstrated using our positive control, *Anolis*. As would be expected, *Anolis* had extremely high enrichment, recovery, and completeness when assembled with BWA against the *Anolis* reference (72%, 99%, and 95%, respectively; Table 2). However, when *Anolis* was assembled against the snake reference a ~27% reduction in both enrichment and completeness occurred (when only genes present in both references were compared). This demonstrates that the use of proper reference sequences is essential for the recovery of complete genes. Furthermore, these results imply that the targeted capture method employed here is highly tolerant to divergence, much more so than current guided-assembly programs.

### Different assemblers performed best on similar and divergent reads

Several different assembly programs were used and their effectiveness evaluated: BWA-MEM (Li 2013), NGM (Sedlazeck et al. 2013), Stampy (Lunter and Goodson 2011), and Bowtie2 (Langmead and Salzberg 2012) (Table 4, Tables S3–S6). BWA and Bowtie2 are both Burrows Wheeler transform-based methods and were designed for assembly of reads to their reference genome, while NGM and Stampy are hash-based methods and were designed for assembly to moderately divergent or polymorphic reference genomes. Thus, we are using these methods in unorthodox ways not only in assembly to divergent species, but also in assembly to complete coding regions rather than whole genomes.

**Table 4.**
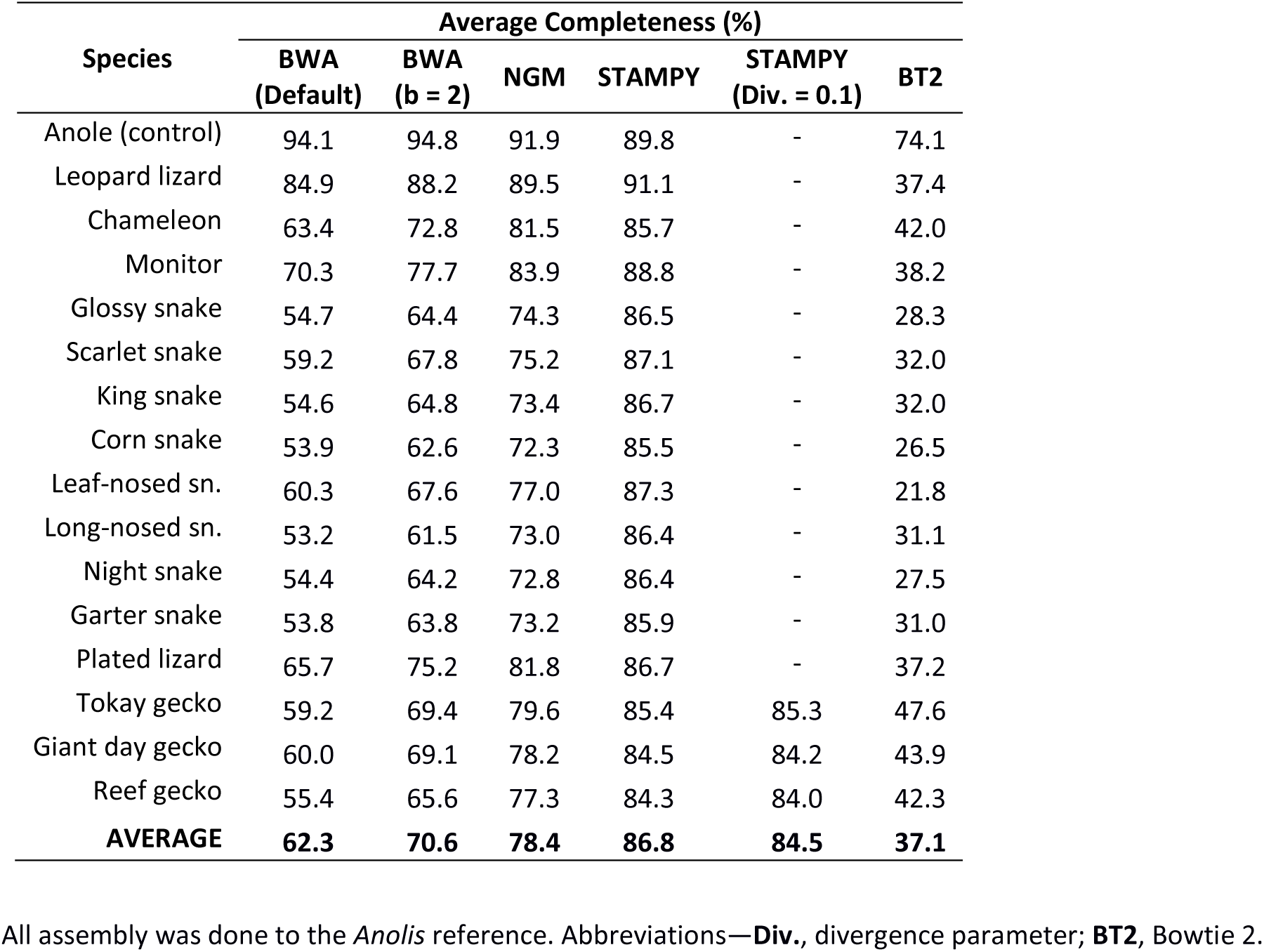
Comparison of average completeness of recovered coding regions obtained using different assemblers.

Initially we implemented BWA under default parameters, but found these to be too restrictive for divergent assembly (Table 4), so we relaxed the mismatch penalty to facilitate cross-species assembly (see Materials and Methods). This improved divergent capture (~10% increase in average completeness), while giving similar results for the positive control (Table 4). Similarly, for the more divergent species we found a 10% increase in going from BWA to NGM and another 10% from NGM to Stampy (Table 4). For the geckos, we found a 17% increase in completeness going from BWA to Stampy. However, this increase in completeness was almost removed when the gecko reference (rather than the *Anolis* reference) was used (2% increase in completeness). For *Anolis*, Stampy actually performed more poorly than BWA, as did NGM (Table 4).

Overall, we found that Stampy performed best when assembling reads to more divergent references. However, this increased ability for divergent assembly appears to have come at the cost of completeness and accuracy in some cases. We found that Stampy incorporated more ambiguous bases than BWA and occasionally resulted in unambiguous differences. This was most apparent in genes with lower completeness, whereas genes with high completeness were most often found to have identical sequences between BWA and Stampy (and NGM). This is not surprising given genes with lower completeness likely had higher divergence, and had much lower depth of coverage (presumably due to lower enrichment), which makes it more difficult to reliably map reads and more likely that incorrect read placements would be accepted. As a result, we tended to prefer BWA assembled sequences to Stampy, but when the divergence was high Stampy was able to capture much more of the gene. In some cases it was possible to combine BWA and Stampy sequences, using BWA to resolve ambiguities and differences and Stampy to fill in missing, presumably divergent regions.

In addition to running Stampy under default settings, we also adjusted the substitution rate option, which should improve mapping of divergent reads. We changed the default rate of 0.001 to 0.1, which corresponds to an expected divergence of 10% and ran this for the most divergent species, the geckos. However, changing this parameter did not have a positive effect on the sequences recovered and perhaps resulted in slightly less completeness (Table 4).

Compared to BWA and Stampy, NGM resulted in intermediate completeness when reads were aligned to a divergent reference (Table 4). When reads were aligned to a more similar reference (e.g., *Anolis* to *Anolis* or a colubrid to the snake reference), however, NGM performed worse than BWA. As such, we did not find NGM to be particularly useful for assembly of either divergent or non-divergent reads.

Lastly, we implemented Bowtie 2, which has recently been used to assemble target enrichment sequencing reads from cichlid fishes (Ilves and Lopez-Fernandez 2014). We ran Bowtie 2 in our assembly and analysis pipeline using the ‘very-sensitive’ preset, which was used by Ilves and Lopez-Fernandez (2014), against both the *Anolis* and snake references. We found that Bowtie 2 performed worse than BWA in all respects (Table 4, Tables S3–6). For example, with the positive control we found a 29% reduction in enrichment and a 21% reduction in completeness. This is a surprising result, but may be due to issues associated with assembly to coding sequences rather than a complete (mammalian-size) genome, which Bowtie 2 was designed to assemble against. Overall, these results highlight the important fact that no single assembler will be best in all situations.

### Increased probe diversity and tiling substantially increase gene recovery and completeness

To evaluate the effect of increased probe diversity and higher levels of tiling we increased both for a small subset of genes. For the visual opsins we included probe sequence from nine different species (including a colubrid snake and a gecko) and doubled the amount of tiling to 20X. Because we had these additional sequences, we also used additional references to assemble this subset of genes. The result was near complete recovery of the visual opsin genes, with the exception of those genes that appear to have been lost in particular lineages (Table S7). These results suggest that when fully complete coding regions are required both the number of probe and reference sequences and, presumably to a lesser extent, the amount of tiling should be increased. Unfortunately the experimental design did not allow us to differentiate between the effects of the number of probe/reference sequences and the amount of tiling, and this will need to be evaluated in a future study. It seems likely that the most important factor in the recovery of complete coding regions is the availability of probe and reference sequences that are as similar as possible to the target, but increased tiling may provide additional benefits when this is not possible.

### Short exons had only a small effect on completeness of recovered genes

One of the largest drawbacks of targeted capture may be difficulty in capturing short targets. This is especially true when targeted exons are shorter than the probe length, which in our case was 120 bp. Including flanking intron sequence to make up the remaining sequence is ideal when doing targeted resequencing, but for cross-species capture is more problematic due to higher sequence divergence among introns. Instead, exons less than 120 bp were padded with non-homologous sequence. To determine what effect this had on the capture of short exons we compared completeness between genes that did not have exons less than 120 bp with those that did. We found, with the non-parametric Mann-Whitney test, that genes with exons less than 120 bp had significantly lower completeness; however, this difference was small with average completeness only being 4% less (Table S8). This difference was similar when the cutoff was set to 100 bp and 50 bp (Table S8).

While significant, the difference in completeness was quite small and many genes with short exons were captured with high (or full) completeness, including the regions comprised from the short exons. For example, *ABCA4* which has 14 exons under 100 bp in length with our *Anolis* probes was captured with 98.3% completeness in *Anolis* and 87.7% on average across the 16 species. The area of the sequence that was not captured in *Anolis* corresponded to a section of sequence where the probe sequence extracted from ENSEMBL and the current predicted sequence on NCBI disagreed. The lack of capture of this area appears to be due to incorrect probe sequence rather than a short exon. *USH1C*, which also had 14 exons under 100 bp, as well as four under 50 bp, was captured at 98.5% in *Anolis* and 90.9% overall. Similarly, the areas not captured in *Anolis* were portions of the sequence that disagreed between the ENSEMBL exons we used to design the probes and the (most recent) NCBI predicted CDS. For our method, short exons do not appear to be a major determinant of the completeness of recovered exons and thus do not represent a substantial obstacle.

### Incomplete and erroneous probe sequences caused substantial reductions in gene completeness

As noted above, one reason that sequences may not be captured is if they were missing from the probe sequence, either due to an incomplete or erroneous sequence. The probe sequences we used were based on the ENSEMBL and NCBI gene predictions available at the time, but the gene predictions have been updated considerably since the probes were developed, especially following the reannotation of the *Anolis* genome (Eckalbar et al. 2013). Though we manually curated the set of sequences and corrected errors when possible using a multiple sequence alignment, it was not possible to fix all of the errors. The most common incongruence between the sequences used for probe design and the updated gene predictions was in the prediction of the first and last exon of the gene. In many cases the first or last exon was incorrectly predicted, but in some cases may have represented an alternate transcript variant. In other cases the updated sequence was actually incorrect based on a multiple sequence alignment (e.g., *CNGA3, GNB5*). More rarely other sections of the sequence would be missing or had insertions (presumably intronic sequence), but these differences were much easier to identify and fix.

When the probe sequence was incomplete or erroneous that part of the sequence was often not captured. This resulted in an overall lower completeness for those genes with missing (72%) or incomplete (76%) *Anolis* probes or references compared to those with complete probes and references (93%, Table S4). This difference was not as substantial as we expected, likely because the additional reference sequences we used allowed the sequence to still be captured in some cases. This is evident from four genes (CNGA1, CRYZL1, ENO1, PGK1) that lacked an *Anolis* probe (but had an *Anolis* reference sequence for assembly) and still had an average completeness of 82% despite being captured with only turtle or chicken probes.

### Completeness of recovered genes decreased with increasing sequence divergence

One of the most important aspects to consider when designing a cross-species sequence capture experiment is the level of sequence divergence that can be tolerated. Many previous methods have targeted sequences with low divergence and/or only closely related species (Mason et al. 2011; Bi et al. 2012; Crawford et al. 2012; Faircloth et al. 2012; Lemmon et al. 2012; McCormack et al. 2012; Ilves and Lopez-Fernandez 2014). Instead, we have targeted a broad range of both sequence divergence and relatedness including species that are up to 200 myr divergent from the closest probe sequences used. In order to evaluate the effect sequence divergence had on the recovery of the coding regions, we developed two approaches for independently calculating divergence. This was necessary because directly measuring divergence between the captured data and the reference would highly bias the results towards the captured data. First, we calculated an average sequence similarity between *Anolis* (the reference) and each of the 15 other species using three independently sequenced genes as a proxy (Table S9). This metric provided a rough estimate of average sequence similarity between each of the species and the probe/reference sequences. However, it was not possible to separate the effects of the hybrid capture and the guided assembly on the completeness of the recovered genes with this approach. To more directly measure the effect of divergence on the hybrid capture we utilized the *Gekko* reference, which allowed us to directly and independently calculate sequence identity between the *Anolis* and *Gekko* sequences. We then compared similarity to completeness calculated through BWA assembly against the *Gekko* reference, which removed the effect of the cross-species assembly and allowed a more direct evaluation of the effect of sequence divergence on hybrid capture (Table S10).

Our measure of species-level sequence similarity revealed a strong correlation with completeness (Fig. 3A, Table S9; *r* = 0.95 *p* < 0.001). The colubrid snakes, which showed the highest level of sequence divergence (despite being more closely related to *Anolis* than the geckos) also showed the lowest completeness. However, despite having average pairwise identities to *Anolis* below 80%, average completeness was above 85%. Variation within the group may represent specific differences in the sequences or variation in DNA or library quality (e.g., the low completeness of *Rhinocheilus*). The lowered completeness of *Sphaerodactylus* is likely due at least in part to divergence from the *Gekko* sequences used as a reference for assembly. Rather than implying a strict relationship between divergence and completeness, these results highlight multiple factors that correlate with evolutionary divergence, which are likely to affect both the hybrid capture and the computational assembly. Generally, these results imply that acceptable levels of completeness can be expected down to 75% sequence similarity when a reasonably close reference is available for assembly. At 90% similarity most genes can be expected to be nearly complete.

**Figure 3.**
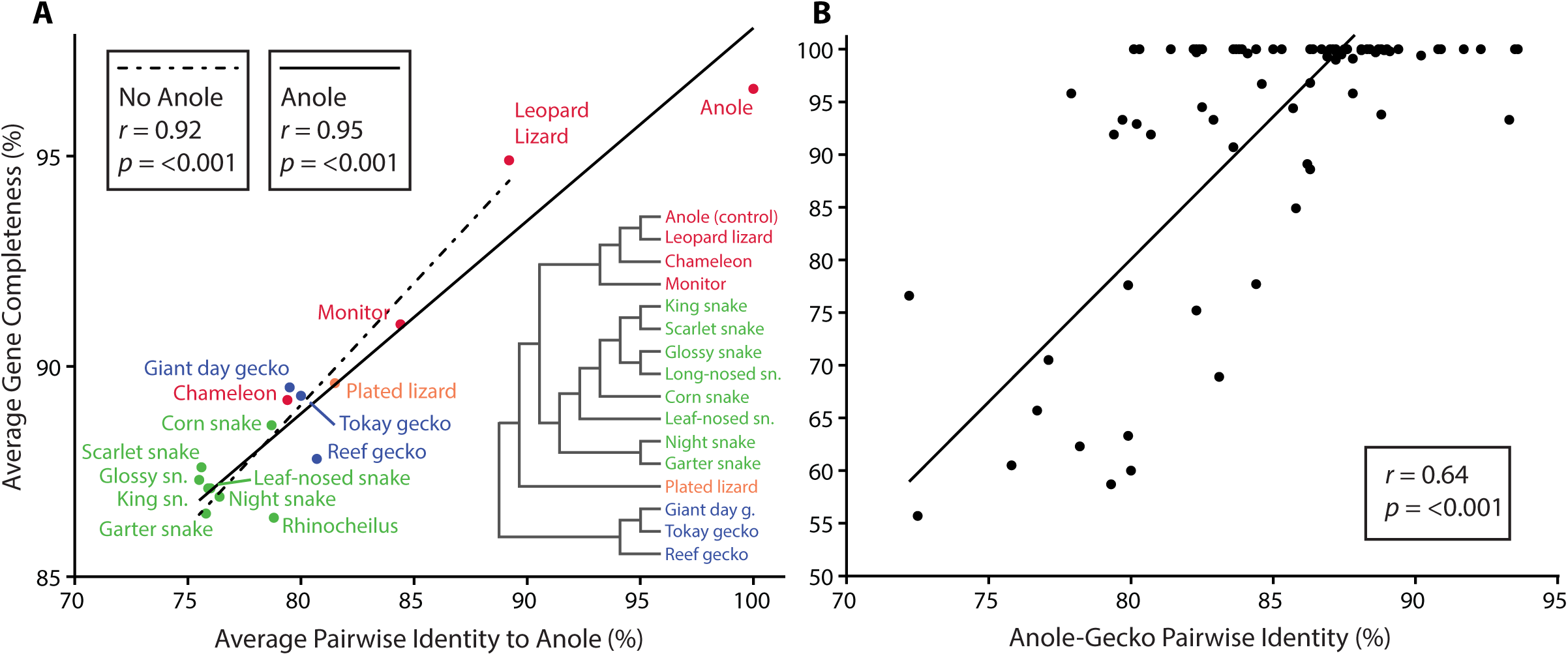
Analyses of the effect of divergence on completeness of recovered coding regions. **A**, average gene completeness for each species compared to average pairwise sequence identity to anole demonstrating the strong correlation between sequence identity and the completeness of genes recovered. Average completeness was calculated as the average across each gene recovered using the best assembly method and reference for each gene. Pairwise identity was calculated between each species and anole for a set of representative genes obtained independently for each species. The reduced major axis regression lines are shown both including and excluding *Anolis* in the regression. **B**, completeness of each gene captured in *Gekko* compared to the pairwise identity between *Anolis* and *Gekko* for those genes. Completeness was calculated for genes with complete probe and reference sequences in *Anolis* that were also found in the *de novo* transcriptome assembly of *Gekko* and are the values for BWA assembly against the *Gekko* reference. Pairwise identity was calculated between the *de novo* assembled *Gekko* coding sequences and the *Anolis* reference sequences thus removing bias from the guided assembly approach.

The level of sequence divergence tolerated with our approach compares favourably with the transcriptome-based exon capture method of Portik et al. (2016). Using a similar approach, the authors compared average pairwise divergence from the probe design species to completeness. Similarly, the authors found a strong linear relationship between divergence and completeness, but with a stronger slope and lower completeness at equivalent levels of divergence (Portik et al. 2016). For example completeness at 10% divergence (90% similarity) was only that at ~70% compared to over 90% in our study. At the other end completeness at 20% divergence (80% similarity) was ~20% compared to over 85% (Portik et al. 2016).

The *Gekko*-specific metric, which allowed us to more directly compare the effects of sequence divergence on the completeness of recovered genes, showed a weaker, but still highly significant, correlation (Fig. 3B, *r* = 0.64, *p* < 0.001). This suggests that, as noted earlier, sequence divergence has a strong negative affect on the performance of the guided assembly. When the effect of the cross-species assembly is removed (by assembling *Gekko* reads to a *Gekko* reference), the correlation weakens, but still accounts for the majority of variation in completeness. Thus, at the individual gene level, aspects other than just overall sequence similarity can have a large effect on the performance of the hybrid capture. As noted above, the accuracy of the probe sequence can have a large affect, but other aspects, including overall similarity in gene structure, concentration of differences, secondary structure, and GC content, may also contribute significantly. The large amount of variation in completeness is exemplified by the large range of sequence identities, from 80–94%, for those genes that were recovered with 100% completeness. When these genes were removed, the correlation was strengthened somewhat (*r* = 0.71), but the remaining comparisons still show a large amount of variation. These results demonstrate that sequence similarity down to 78% between the probe sequence and the target can still result in the capture and enrichment of nearly complete (95%+) coding sequences.

### Targeted capture performed similarly or better, and cost as little or less, than RNA-Seq and WGS

To demonstrate the usefulness of our method for sequencing complete coding sequences in comparison to other approaches, we assembled RNA-seq reads from *Thamnophis sirtalis* eye tissue and previously published whole genome sequencing datasets (Card et al. 2014; Zhang et al. 2014) of various coverage using our guided assembly pipeline. We evaluated the recovery and completeness of the assembled sequences, as well as the costs of sequencing, in comparison to our targeted capture approach. We found that the RNA-Seq dataset recovered substantially fewer genes, but that the completeness of genes that were recovered was similar to that of the capture data in general, and slightly higher than the capture for *Thamnophis* specifically (Table 5, Table S11). The reduced number of genes recovered was due primarily to the fact that RNA-Seq was performed on eye tissue and not all of the 160 genes that were targeted are expressed in the eye, despite the focus on visual genes in our probe set. This highlights one of the main drawbacks of RNA-Seq, but could be overcome if the genes of interest were all sufficiently expressed in a single tissue type or by pooling RNA extracted from multiple tissue types, although the second option may require additional sequencing depth. Additionally, genes may not have been recovered if they were not expressed at sufficient levels to be captured at the sequenced coverage level. Sequencing rare transcripts will be more difficult (i.e., require higher coverage) with RNA-Seq, but this drawback does not apply to targeted capture or other genomic approaches.

**Table 5.** Comparison of the performance of the assembly and annotation pipeline on RNA-Seq and whole genome data with the targeted capture approach.

| Sequencing Method | Species | Number QC-passed Reads | Percent Reads Mapped | Genes Recovered | Average Completeness |
| --- | --- | --- | --- | --- | --- |
| Targeted capture | Average of 16 | 8,567,574 | 51.6% | 147 (92%) | 70.6% |
| RNA-Seq | *Thamnophis sirtalis* | 29,843,391 | 4.29% | 112 (70%) | 73.5% |
| WGS | *Centrocercus minimus* (SRR1166456) | 26,817,128 | 0.05% | 40 (26%) / 106 (68%) | 12.6% / 30.8% |
| WGS | *Nucifraga columbiana* (SRR1166560) | 43,587,682 | 0.05% | 42 (27%) / 135 (87%) | 12.6% / 40.4% |
| WGS | *Tyto alba* (SRR959575) | 184,107,287 | 0.02% | 31 (20%) / 85 (55%) | 26.8% / 39.0% |
| WGS | *Struthio camelus* (SRR950910) | 219,605,123 | 0.02% | 122 (79%) / 151 (97%) | 48.6% / 76.2% |
| WGS | *Calypte anna* (SRR943144) | 265,237,073 | 0.10% | 155 (100%) | 82.9% / 88.2% |

Data shown is for assembly with BWA against the *Anolis* (targeted capture, RNA-Seq) or *Gallus* reference (WGS). Data for individual genes is shown in Tables S11 and S12. Since the whole genome sequencing (WGS) suffered from low mapping rates (due to no enrichment) we additionally relaxed the requirement in our assembly pipeline to only require the minimum (default) level of coverage of 3X, which is shown after the slash (/). A sequence was considered recovered if it had a minimum of 5% completeness and could be identified via BLAST. Completeness was calculated relative to the reference sequence for those genes identified as recovered.

While the completeness levels were similar between reference guided RNA-Seq and targeted capture, we also compared results for *de novo* assembly of the transcriptome as this would be the typical procedure with a species that lacks a reference genome. With *de novo* assembly we recovered 77.5% of the 160 genes with an average completeness of 91.5% for *Thamnophis* (Table S11). However, our guided assembly pipeline is highly stringent in that it requires a minimum of 10X coverage for a base to be called in the consensus. When we relaxed this to the default minimum of 3X we found an increase in recovery and completeness that slightly exceeded the *de novo* assembly.

We also compared our targeted capture method against non-enriched whole genome sequencing (WGS). We selected four previously sequenced datasets containing an increasing number of reads in order to determine what sequence coverage was necessary to overcome the lack of enrichment. Using our same assembly and analyses pipeline, we found that only at the highest sequencing coverage tested (265 million reads) was recovery and completeness satisfactory (Table 5, Table S12). However, when we relaxed the stringent 10X coverage requirement to the default minimum of 3X we obtained a marked increase in both recovery and completeness, but this was still well below that found for the sequence capture experiment for the low and medium coverage tests (26–184 million reads).

In terms of the costs of producing the data, RNA-Seq and targeted capture are similar, whereas whole genome sequencing was substantially more expensive. A single targeted capture sample (for a run with 16 samples total) and ~30 million reads of RNA-Seq (approximately one sixth of a HiSeq lane) cost essentially the same (Table 6, Table S13). Sequence capture further excels due to its scalability to larger numbers of samples, which at 96 samples would have reduced the cost by almost 20% per sample. When smaller numbers of samples are required, and RNA appropriate tissue is available, RNA-Seq would be the preferred method due to the time investment involved in developing a set of probe sequences and the relative ease of *de novo* assembly. For genome sequencing, even the low coverage genomes cost more than targeted capture (per sample), with the higher coverage genomes costing almost four times as much. Despite the substantially higher cost, WGS may be desirable if there is a very large number of genes of interest and/or if there is only a small number of species to be sequenced. RNA-Seq may still be preferable in these cases if fresh tissue is available. Otherwise, sequencing a species on the equivalent of a single HiSeq lane (our highest coverage tested, but still much lower than needed to *de novo* assemble a complete genome) may be a useful way to obtain many nearly complete coding sequences. A cross-species reference-guided genome assembly approach, as proposed by Card et al. (2014), may be able to recover more complete genes from lower coverage genomes as well.

**Table 6.** Cost comparison of targeted capture, RNA-Seq, and whole genome sequencing experiments.

| Per Sample Costs | Extraction | Kit & Reagents | Library Prep | Fraction of HiSeq Lane | Cost of Sequencing | Total | Relative Cost |
| --- | --- | --- | --- | --- | --- | --- | --- |
| Targeted Capture 16 | \$3.81 | \$482.74 | \$100.00 | 1/32 | \$65.63 | \$652 | 100.0% |
| Targeted Capture 96 | \$3.81 | \$367.79 | \$100.00 | 1/32 | \$65.63 | \$537 | 82.4% |
| RNA-Seq | \$14.45 | - | \$300.00 | 1/6 | \$350.00 | \$664 | 101.9% |
| WGS | \$3.81 | - | \$251.00 | 1/7 | \$300.00 | \$555 | 85.1% |
| WGS | \$3.81 | - | \$251.00 | 1/5 | \$420.00 | \$675 | 103.5% |
| WGS | \$3.81 | - | \$251.00 | 1/4 | \$525.00 | \$780 | 119.6% |
| WGS | \$3.81 | - | \$251.00 | 3/4 | \$1,575.00 | \$1,830 | 280.6% |
| WGS | \$3.81 | - | \$251.00 | 1 | \$2,100.00 | \$2,356 | 361.2% |

### Captured phylogenetic markers produced an accurate species tree

The most common application of cross-species sequence capture is for phylogenetic analysis, and our method can also be applied for this purpose. We targeted 23 genes previously used as phylogenetic markers in reptile and squamate phylogenetic studies. Of those, 16 were over 80% complete for all species (Table S14) and so were used to construct a multigene phylogenetic species tree using MrBayes. The resulting tree was highly supported and closely matched a recent multigene squamate tree (Figure 4; Pyron et al. (2013)). The only topological difference was in the position of *Pantherophis*. which is unsurprising due to the extremely short branch lengths in this portion of the tree. These results demonstrate that it is possible to get high quality data for phylogenetic reconstruction that can also be used to address a variety of other research questions.

**Figure 4.**
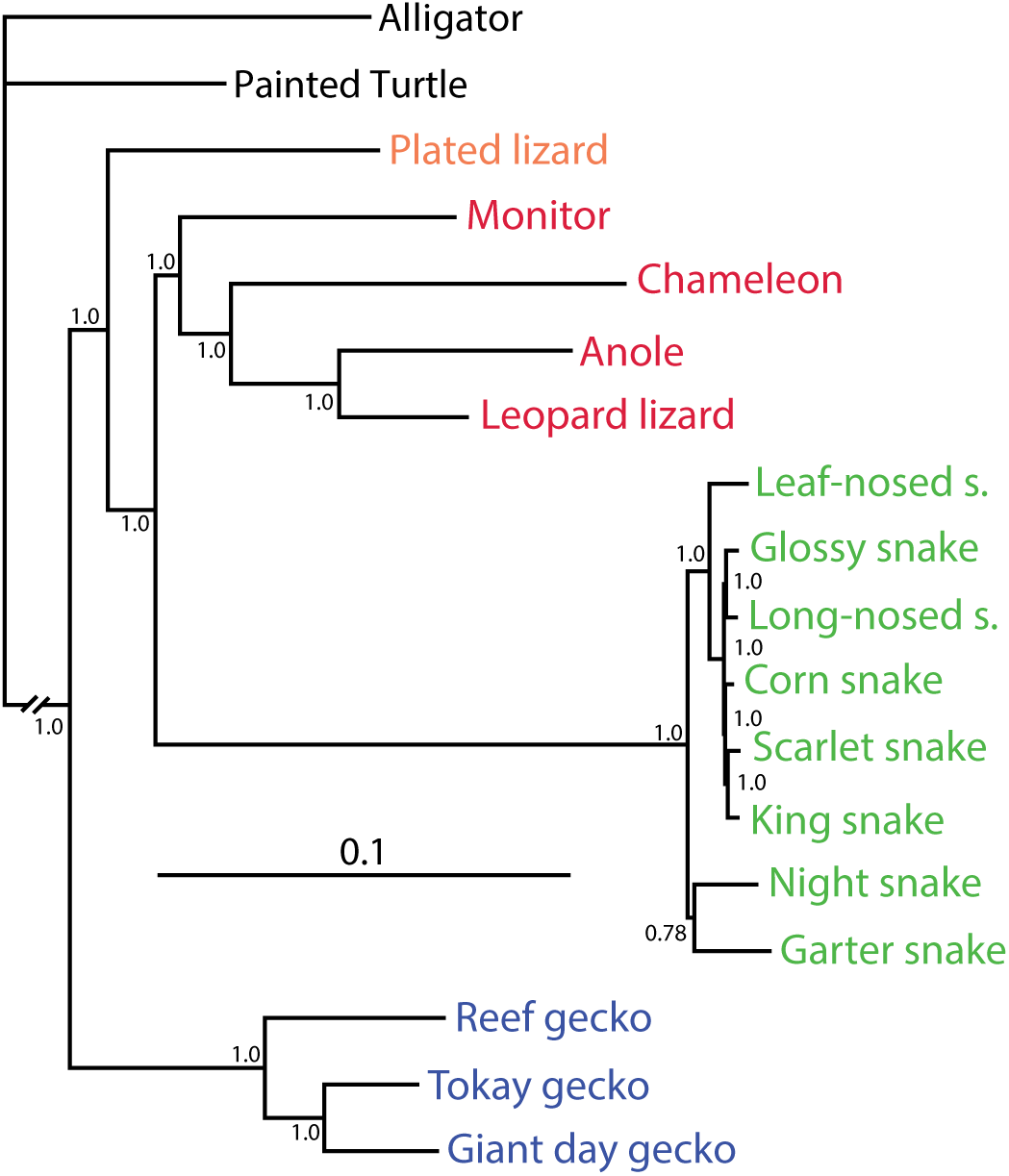
Bayesian multigene phylogeny of the 16 species with two outgroups. A total of 16 phylogenetic marker genes were used. The topology agrees strongly with the multigene phylogeny of Pyron et al. (2013) differing only in the placement of the corn snake. Posterior probability support is shown at each node.

### Conclusions

Overall, the cross-species targeted capture method proposed here was highly successful in recovering the 160 genes of interest with high completeness over large evolutionary distances (up to 200 myr of divergence). Our use of complete coding regions from specific genes of interest allowed us to focus on aspects of organismal physiology producing data useful for both phylogenetics and studies of molecular evolution and function. Recovery of more divergent sequences was lower, but this was primarily due to the difficulty of cross-species guided assembly rather than a failure of the hybrid enrichment. This issue was partly overcome through the use of additional reference sequences obtained from whole genome data. Since the hybrid enrichment appears to have been highly robust to sequence divergence, development of a *de novo* assembly pipeline that removes the reliance on cross-species assembly is a promising avenue for future research. A *de novo* approach, however, is not trivial and our preliminary attempts have produced results worse than or on par with the guided approach developed here. Differences in assembly methods between cross-species enrichment approaches likely accounts for a large amount of the variation in the quality of divergent capture and needs to be further evaluated. Because our results show substantial increased recovery when additional probe sequences are included, adopting a transcriptome-based targeted capture approach similar to that proposed by Bi et al. (2012) and Portik et al. (2016), where transcriptomes from one or more species are first sequenced and *de novo* assembled and then used to design a set of probes for hybrid capture, may be highly beneficial. Modifications to the targeted capture protocol, as well as introduction of a second round of capture, as implemented by Li et al. (2013) may further extend the ability to capture divergent sequences. The data produced by our method was more than sufficient to produce a robust phylogenetic tree and will be used for future molecular evolutionary and functional studies. While we initially targeted a modest number of genes and species the method is easily scalable to much large numbers of both, which will further increase its efficiency and per sample cost effectiveness. The cross-species targeted capture method developed here will enable the study of a variety of evolutionary questions in virtually any set of genes of interest across divergent groups of species.

## Materials and Methods

### Probe Design

As a proof-of-concept for this method we targeted 166 visual, housekeeping, and phylogenetic marker genes. This set of genes included nearly all genes known to function in the phototransduction and visual cycles, as well as genes involved in photoreceptor development and maintenance, non-visual opsins, and lens crystallins (Table S1). This set of genes was collected, in part, from proteome and transcriptome papers of photoreceptor outer segments and retinas (Schulz et al. 2004; Kwok et al. 2008). Phylogenetic markers were selected from reptile and squamate phylogenetic studies (e.g., Harshman et al. 2003; Iwabe et al. 2005; Vidal and Hedges 2005; McAliley et al. 2006; Hugall et al. 2007; Barley et al. 2010) and housekeeping genes from the list produced by She et al. (2009). In order to promote cross-species hybridization, probes were designed from a representative set of taxa that have complete genomes spanning reptilian phylogenetic diversity, including *Anolis* (lizard), *Pelodiscus*/*Chrysemys* (turtle), and *Gallus* (bird), following Lemmon et al. (2012). This ensured that a range of sequence variation was present in the probe sequences to promote hybridization with divergent sequences. For each of the 166 targeted genes, we obtained mRNA or predicted mRNA sequences from Genbank and coding sequences (CDS) and individual exon sequences from ENSEMBL, as available for each of the probe taxa. When exon sequences were not available on ENSEMBL we attempted to obtain them through direct BLAST searches of the genomes. The individual exons were aligned to the complete coding and/or mRNA sequence using custom scripts and manually inspected and corrected as necessary. Sequences from all probe taxa were aligned together, which allowed intronic and UTR sequences present in the exon annotation to be identified and removed. We found such contaminating sequences to be common in the exon sequences obtained from ENSEMBL as intron-exon boundaries and start and stop codons were often misidentified. This step also allowed us to manually verify the annotation of each gene and exon. All 166 genes were not present in the reference genomes and thus some genes are only represented by sequences from only one or two species and in a few cases substitutes were used (Table S1). If sequences for both *Pelodiscus* and *Chrysemys* were available only the longer sequence was kept. If they were of the same length and overall quality the *Pelodiscus* sequence was given preference for consistency, as it had the more complete genome in general. Once exons were validated, the mRNA and CDS sequences were removed and only the exons retained. For the opsin genes, we added additional sequences from lizards, snakes, and alligator (as available) to the probe sequences. These sequences were manually broken into their constituent exons based on the multiple sequence alignment including the exons from ENSEMBL for *Anolis*, *Pelodiscus*/*Chrysemys*, and *Gallus*. Additional probe species were added for the opsins in order to evaluate the effect of increased probe diversity on capture efficiency and to ensure complete capture of these genes. Altogether, the process resulted in a total of 3888 exons from which the probes were designed. A complete breakdown of the genes and exons targeted by the probes is available in Tables S1 and S8.

The probes consisted of 120 bp of RNA synthesized by Agilent, which is the median size of protein coding exons in the human genome (Clamp et al. 2007) and because RNA has stronger hybridization with DNA than DNA does. Probes were extensively tiled across the exons (10X coverage, 20X coverage for opsin genes) to increase the likelihood of hybridization of inserts with at least a single probe variant, with the goal of increasing capture of complete exons. Exons that were shorter than the probe length were padded with non-homologous sequence because probe length could not be shorter than 120 bp. The number of probes targeting short exons was boosted in order to normalize coverage of the target region. This resulted in a total of 45,895 probes after tiling and boosting.

### Sample Preparation and Sequencing

To test the method, 16 squamate reptiles were selected that varied in their divergence from *Anolis*, including eight snakes, three geckos, and five other lizards (Table S2). As a positive control, we included *Anolis* (the squamate probe species) in this set of 16 species. This range of species spans much of the diversity of squamates and allowed for evaluation of the efficiency of capture and enrichment at different levels of sequence divergence. Genomic DNA (gDNA) was extracted from the muscle and/or liver samples using the DNeasy Blood and Tissue Kit (Qiagen) following the manufacturer’s protocol. Library creation, hybridization and sequencing were performed according to the Agilent SureSelect protocol at the Centre for Applied Genomics (TCAG; Sick Kids Hospital, Toronto). The 16 samples were sequenced on roughly half a HiSeq (Illumina) lane (approximately 1/32 of a lane per sample).

### Reference File Creation for Guided Assembly

To facilitate assembly across divergent species, and evaluate the effect of the computational assembly on gene recovery, several different sets of reference files were generated that differed in the primary species used to build the reference. These were an *Anolis*, a snake, and a *Gekko* reference, as well as several additional small references targeting just the visual opsin genes. The *Anolis* reference was constructed using the available *Anolis* sequences from Genbank for the 166 targeted genes. Due to improvements to the *Anolis* genome and its associated gene predictions that occurred after building the probe set, the *Anolis* sequences present in the reference are not necessarily the same as those used to construct the probes. The updated sequences should represent more accurate predictions and thus were used in most cases. In some cases this meant inclusion of an *Anolis* sequence in the reference that was not present in the probe set. If a sequence still could not be found from *Anolis* the next most closely related sequence was used (*Python, Pelodiscus, Alligator, Gallus*). In some cases when only a partial *Anolis* sequence was obtainable the missing portion was added from *Python*. After an initial survey using this reference six sequences were removed. Two genes (ALB, *SLC24A1*) were found to have been ancestrally lost in squamates (not present in any squamate genome or any of the 16 species sequenced in this study). One was identified to be a lineage-specific duplication in some birds (*CRYD2*). The probe for one gene, *STRA6*, was found to lack any homology with other *STRA6* sequences and thus was not successful at capture. Additionally, three genes initially included, *UBC, UBB*, and *UBI*. were found to all represent the same gene (which we term *UBC*) and thus were combined. This left a total of 160 genes that made up the *Anolis* reference.

The snake reference was built primarily from sequences obtained from the *Python* and *Thamnophis* genomes and a *de novo* Trinity transcriptome assembly of *Thamnophis*. The *Thamnophis* sequences were preferred over the *Python* as the included snake species are more closely related to *Thamnophis* than *Python*. If only a partial sequence was obtainable, *Anolis* sequence was used to complete it when possible. This resulted in 139 sequences in the snake reference. A third reference was also used and this was based on sequences obtained from the *Gekko japonicus* genome (110 sequences).

### Assembly and Analysis Pipeline

Raw reads were processed by Trimmomatic (Bolger et al. 2014) to remove low quality reads, as well as primer and index contamination under default settings. A complete pipeline was developed for the assembly and analysis of trimmed reads. First, reads were assembled using one of three methods: BWA-MEM (Li 2013), NGM (Sedlazeck et al. 2013), or Stampy (Lunter and Goodson 2011). BWA-MEM is the most conservative method in terms of tolerating mismatches between the reads and the reference, but is also the most accurate, whereas NGM and Stampy both tolerate more mismatches, but at the cost of some accuracy (Lunter and Goodson 2011; Li 2013; Sedlazeck et al. 2013; Turki and Roshan 2014). However, the benefit of allowing more mismatches in assembling reads to divergent reference sequences may outweigh a small reduction in accuracy. BWA-MEM was first run under default parameters, but assembly was found to suffer when applied across species. To address this, we reduced the mismatch penalty from the default of 4 to 2 (-B 2) and used this for subsequent analysis. NGM was run under default parameters. Stampy was also run under default parameters, but a subset of analyses were run to test the effect of changing the substitution rate parameter. Since Bowtie 2 (Langmead and Salzberg 2012) has been used recently to assemble targeted capture data (Ilves and Lopez-Fernandez 2014) we additionally implemented Bowtie 2 under the ‘very sensitive’ preset used by Ilves and Lopez-Fernandez (2014) in our analysis pipeline using the *Anolis* and snake references.

Consensus sequences were called using the mpileup-bcf-vcfutils pipeline of Samtools (Li et al. 2009) with a minimum sequence and mapping quality score of 20 (-Q 20 and -q 20) and a minimum depth of coverage of 10 (-d 10). Additionally the parameter ‘l’ was set to 1 in vcfutils, which reduced the number of bases surrounding an indel that were replaced with ‘N’s to one. Note that while the probes were targeted to individual exons the assembly was done against complete coding regions. The consensus sequences generated are the assembled coding region of the captured gene. After removing lowercase letters from the consensus sequence (which signify bases that did not meet the quality and depth of coverage standards) the completeness of the recovered coding region, relative to the reference sequence, was calculated using custom scripts. Consensus sequences were annotated by BLAST to identify the recovered gene in comparison to the gene targeted by the reference.

Completeness calculations and BLAST annotations were manually verified for each gene. Genes were considered recovered when they had at least 5% completeness and the BLAST annotations matched the targeted reference sequence. Where BLAST annotations were ambiguous, simple maximum likelihood gene trees were inferred using either PhyML (Guindon et al. 2010) or MEGA (Tamura et al. 2011b) to verify sequence identities. Sequences that did not meet these criteria were removed and not used for further comparative analyses.

### Method Analysis and Evaluation

Completeness of the recovered coding regions was compared across the different reference sets and assembly methods. In addition to completeness, we also compared the enrichment efficiency using the simple proxy of the percentage of reads that mapped to the reference. In order to evaluate the effect of sequence divergence on the recovery of the gene, a proxy for average sequence divergence between *Anolis* and each of the other 15 taxa was calculated. To avoid biasing the results, divergences could not be calculated based on the recovered sequences. Instead, genes that were independently sequenced and available on Genbank for each of the 16 species were needed. Six candidate genes were identified: *BDNF, MOS, NTF3, RAG1, R35/GPR149*, and *ZEB2*. To increase sample size, sequences from different species within the same genus were included. Pairwise identity was calculated between the sequence for *Anolis* and each of the 15 other species using PRANK (Loytynoja and Goldman 2005) to align the sequences followed by USEARCH to calculate a distance matrix (Edgar 2010). Three of the six genes (*MOS, NTF3*, and *R35*) had almost complete taxon coverage and similar average identities with the inclusion of additional species from the same genera. Comparison of identities between multiple species in the same genera revealed very little variation. As such, the average of these three genes was used as a proxy for average sequence identity between the species.

In addition to estimating sequence identity between species, we also calculated levels of sequence identity of the individual genes by utilizing the *Gekko* reference in a more specific, but also more robust, comparison. Pairwise sequence identity was calculated between the *Anolis* and *Gekko* reference sequences (obtained from the genome and thus independent from the target enrichment sequences) for each of the genes present and complete in both species. We compared these sequence identities to the completeness of the recovered coding regions obtained from assembly of the *Gekko* targeted capture reads assembled against the *Gekko* reference. This approach removed the effect of the cross-species assembly, enabling evaluation of the targeted capture efficiency directly.

To compare the effect of increased probe diversity and tiling, we compared the completeness of the visual opsin genes to the overall average. We also investigated the effect of short exons on gene completeness. Because exons shorter than 120 bp were padded with non-homologous sequence, and necessarily could not be tiled, we expected a reduction in the recovery of these exons. To evaluate this, we compared completeness of genes with no exons under 120, 100, or 50 bp to those that had one or more exons under these thresholds. Differences between the two groups were evaluated with the non-parametric Mann-Whitney test as the distributions were highly skewed (non-normal).

### Phylogenetic Analysis

To evaluate the usefulness of the recovered data for molecular evolutionary studies, a multi-gene species tree was inferred using the captured phylogenetic marker genes. Only genes that had both complete coding sequences in the probe and reference files and that were at least 80% complete were used. This resulted in the selection of 16 out of the 23 genes we had identified as phylogenetic markers. Sequences for each of these genes were aligned using MUSCLE (Edgar 2004) codon alignment implemented in MEGA (Tamura et al. 2011a) along with outgroup sequences from *Alligator mississippiensis* and *Chrysemys picta*. Individual multiple sequences alignments were concatenated and partitioned into individual genes. The matrix was analyzed using MrBayes (Ronquist et al. 2012) using reversible jump MCMC with a gamma rate and invariant sites parameter (nst=mixed, rates=invgamma), which explores the parameter space for the nucleotide model and the phylogenetic tree simultaneously. The analysis was run for five million generations with a 25% burn-in. Convergence was confirmed by checking that the standard deviations of split frequencies approached zero and that there was no obvious trend in the log likelihood plot.

### Data Availability

Data associated with the manuscript including probe information, custom scripts, and reference files will be available through DRYAD upon publication.

## Supplementary Materials

Supplementary Tables: Tables S1–S14.

## Acknowledgements

We thank Jiayang Wu for improvements to the assembly and analysis pipeline scripts. We would also like to thank Dante Cerrullo and Agilent for their help with probe design and Sergio Pereira and the Centre for Applied Genomics at Sick Kids for their assistance with the targeted capture and sequencing. This work was supported by a National Sciences and Engineering Research Council (NSERC) Discovery grant (BSWC), an Ontario Graduate Scholarship (RKS), and a Vision Science Research Program Scholarship (RKS).

